# BACH1 promotes hepatocellular carcinoma progression by targeting PDP1 towards the PI3K-AKT-mTOR signaling activation

**DOI:** 10.1101/2025.03.18.643938

**Authors:** Qiqi Bu, Xiaojing Qi, Qing Wang, Shujuan Liu, Shanshan Lin, Zhanhang Wang, Yiguo Zhang

**Affiliations:** The Laboratory of Cell Biochemistry and Topogenetic Regulation, College of Bioengineering, Chongqing University, No. 174 Shazheng Street, Shapingba District, Chongqing 400044, China; College of Animal Science and Technology, Northwest A&F University, Yangling, Shaanxi 712100, China; School of Life and Health Sciences, Fuyao University of Science and Technology, No. 104 Wisdom Avenue, Nanyu Town, Minhou County High-Tech District, Fuzhou 350109, Fujian, China

**Keywords:** BACH1, PDP1, PI3K-AKT-mTOR signaling, hepatocellular carcinoma

## Abstract

Hepatocellular carcinoma (HCC) is remaining to be accepted as a major challenge against public health, because of its high incidence and mortality. Herein, we explored underlying molecular mechanisms by which BTB and CNC homology 1 (BACH1) facilitated progression of HCC. Functional experiments indicated that BACH1 promoted the proliferation, clonogenicity, cycle progression of HCC cells, and inhibited apoptosis. Knockout of BACH1 decreased mitochondrial membrane potential, augmented reactive oxygen species (ROS) generation, induced cell apoptosis, and reduced malignant growth of xenograft neoplasms in mice. Mechanistic experiments revealed that BACH1 bound to the ARE sites in the promoter region of pyruvate dehydrogenase phosphatase catalytic subunit 1 (PDP1), so to activate its transcriptional expression and promote cellular energy metabolism. Interestingly, BACH1 enables activation of the cell growth-related TGF-β1/SMAD signaling pathway, which further promoted the malignant proliferation of HCC cells. Of note, PDP1 functions as a tumor promoter, since lowering its expression levels suppressed the clonogenicity in HCC cells. Furthermore, such downregulation in BACH1 or PDP1 could inhibit the PI3K-AKT-mTOR signaling. In conclusion, these reveal that BACH1 manifests a dominant tumor-promoting factor and is hence viewed as a promising therapeutic target of HCC, because it promotes HCC progression by targeting PDP1 towards PI3K-AKT-mTOR pathway activation.

## 1. Introduction

Worldwide, liver cancer is a commonly diagnosed cancer and one of the major causes of cancer-associated deaths, representing a major public health challenge [1]. Among all forms of liver cancer, HCC is both the most frequent and numerous, accounting for about 80% of total cases. Despite significant advances in diagnosis and treatments, morbidity and mortality for HCC remain considerably high[2, 3]. In this case, it has become crucial to elucidate the molecular biological mechanisms of HCC malignant progression and develop targeted drugs.

BACH1, a major member of the cap’n’collar and basic region-leucine zipper (CNC-bZIP) transcription factors family, forms functional heterodimers with small Mafs that binds to the antioxidant response elements (AREs) in promoter regions of target genes to activate or repress its transcription [4, 5]. BACH1 is broadly distributed in a variety of tissues and cells, plays important role in maintaining heme homeostasis and regulating the transcriptional expression of antioxidative genes [6]. In addition, BACH1 is up-regulated in several human cancers, its increased expression is associated to malignant progression of tumors [7-11]. In ovarian cancer, BACH1 increases cyclin D1, p-AKT, p-p70S6K expression to enhance growth of tumor and promotes the cell metastasis through recruitment of HMGA2 [9]. High expression level of BACH1 in pancreatic cancer presages a poor prognostic which facilitates metastasis by enhancing epithelial-mesenchymal transition (EMT) and repressing epithelial genes [8, 12]. BACH1 also contributes to esophageal squamous cell carcinoma (ESCC) progression through activating transcription of genes associated with angiogenesis (vascular endothelial growth factor C, VEGFC) and EMT, including CDH2, SNAI2, VIM [13]. Stabilization of BACH1 by lowering ROS level can stimulate angiogenesis and metastasis of lung cancer [14, 15]. BACH1 functions as a tumor-promoting factor in HCC, which promotes cell proliferation, migration and invasion [11, 16]. Nevertheless, the specific molecular mechanisms whereby BACH1 facilitates HCC progression are still unknown.

Impaired glucose homeostasis is considered to be a risk factor leading to metabolic diseases such as cancers. Pyruvate dehydrogenase complex (PDC), the compound of key metabolic enzymes for glucose oxidation, performs an essential function in glucose metabolism. It mediates conversion of pyruvate oxidative decarboxylation to acetyl-CoA, a critical step in cellular energy metabolism [17, 18]. The activity of PDC is strictly governed by a series of metabolic enzymes in which pyruvate dehydrogenase phosphatase (PDP) catalyzes dephosphorylation and reactivates PDC [19]. PDP1 is one of the two isoforms of PDP and encoded by human pyruvate dehydrogenase phosphatase catalytic subunit 1 (PDP1) gene. Recently, PDP1 has been shown to be involved in the malignant behaviors of cancers, which is upregulated in human pancreatic adenocarcinoma and prostate cancer, promoting cells proliferation and tumors growth [18, 20]. In colorectal cancer (CRC), PDP1 acts as a crucial oncogenic driver and promotes progression of KRAS mutant CRC by activating MAPK signals [21]. However, the function of PDP1 in HCC has not been found. Moreover, whether there is a connection between BACH1 and PDP1 in the regulation of HCC progression remains unexplored.

To address them, we here discovered that knockout of BACH1 reduced mitochondrial membrane potential, induced cell apoptosis and suppressed cell growth related TGF-β1/SMAD signaling pathway, thus inhibiting the malignant proliferation of HCC cells. For another, PDP1 functioned as a tumor-promoter, because the clonogenicity in HCC cells was suppressed through lowering its expression level. BACH1 bound to the ARE sites in the promoter region of PDP1, activated its transcriptional expression and promoted cellular energy metabolism.

In addition, down-regulation of BACH1 or PDP1 could suppress PI3K-AKT-mTOR signaling. Taken together, BACH1 manifests as a dominant tumor-promoting factor which promotes HCC progression via targeting PDP1-PI3K/AKT/mTOR signaling pathway. This study emphasizes that BACH1 has great potential to be an effective molecular target for HCC treatment and prevention.

## 2. Materials and methods

All materials and methods see the Supplementary file.

## 3. Results

### 3.1 Knockout of BACH1 diminished the malignant growth of HCC cells in vivo

To verify the function of BACH1 in HCC and further explore the specific molecular mechanisms, we established two mutant cell lines, named *H-BACH1^−/−^* and *LM3-BACH1^*−/−*^*, using CRISPR/Cas9 for gene editing in HepG2 and HCCLM3 cells, respectively. We first identified their genotypes by genomic DNA sequencing (Figure 1A) and confirmed that BACH1 protein was successfully knocked out by Western blotting (Figure 1, B & C). The qPCR results revealed that mRNA expression levels of BACH1 in two mutant cell lines had no obvious changes compared to control values in wild type cells (Figure 1, D & E). These suggested that in mutant cells, the total mRNA level of BACH1 was unchanged, but its protein translation was missing. Then, the exogenous BACH1 overexpression plasmid was transfected into *H-BACH1^*−/−*^* and *LM3-BACH1^*−/−*^*to determine the molecular weight of BACH1 (92 kDa) and verify its knockout again (Figure 1, F & G). In HepG2 cells, control and stably BACH1-overexpressing cell lines were established with the lentiviral system, called LV-Con and LV-BACH1, and the expression efficiency of BACH1 was further confirmed by qPCR (Figure 1H) and WB (Figure 1I).

**Fig. 1.**
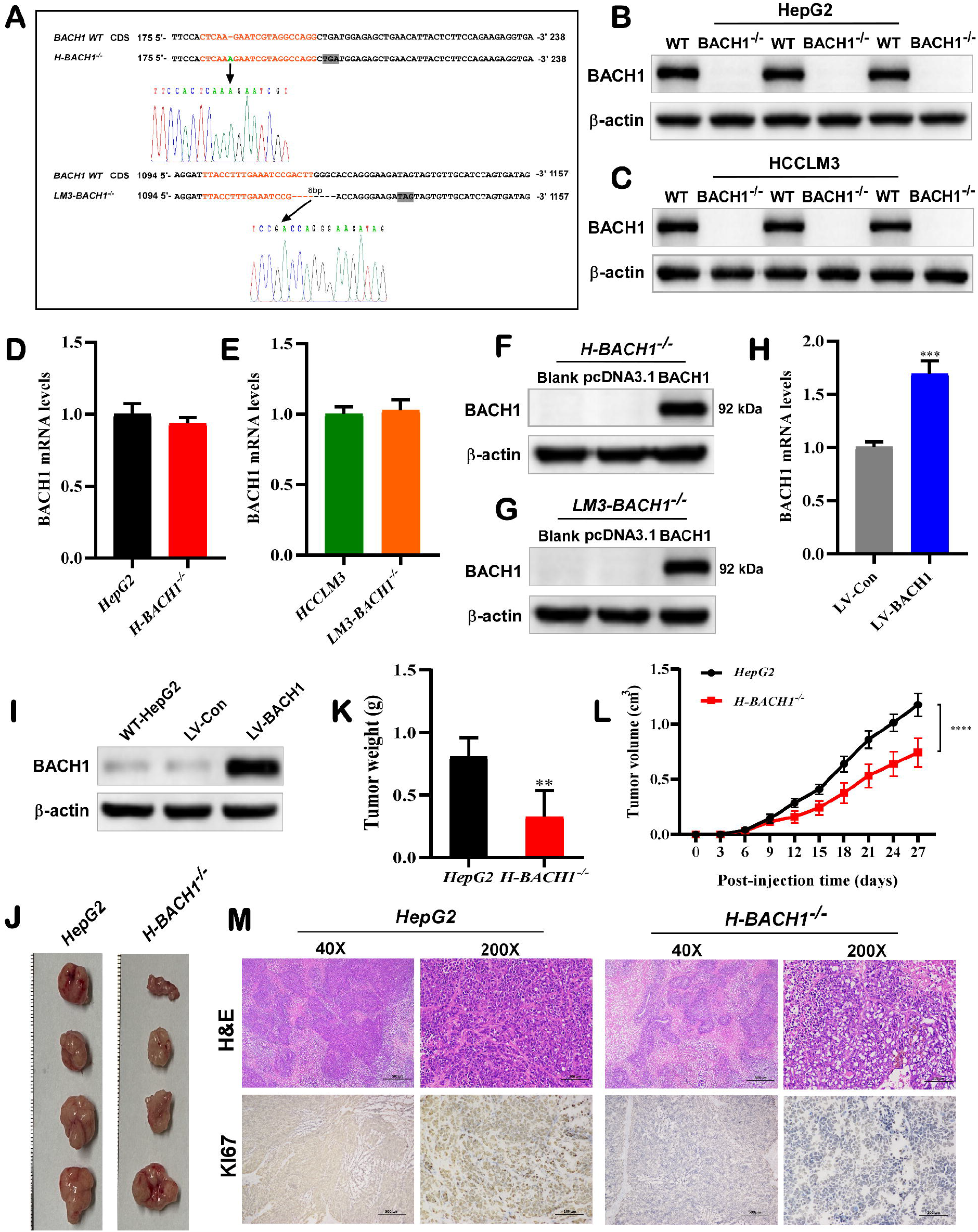
Knockout of BACH1 diminished the malignant growth of HCC cells in vivo. (A) The genomic DNA sequencing results of *H-BACH1*^*−/−*^ and *LM3-BACH1*^*−/−*^ cell lines. (B) The protein expression levels of BACH1 in *HepG2* and *H-BACH1*^*−/−*^ cell lines. (C) The protein expression levels of BACH1 in HCCLM3 and *LM3-BACH1*^*−/−*^ cell lines. (D) The mRNA expression levels of BACH1 in *HepG2* and *H-BACH1*^*−/−*^ cell lines. (E) The mRNA expression levels of BACH1 in *HCCLM3* and *LM3-BACH1*^*−/−*^ cell lines. (F) *H-BACH1*^*−/−*^ cells were transfected with BACH1-expressing constructs, and then the BACH1 protein expression level was examined. (G) *LM3-BACH1*^*−/−*^ cells were transfected with BACH1-expressing constructs, and then the BACH1 protein expression level was examined. (H) The BACH1 mRNA levels were analyzed in the LV-Con and LV-BACH1 cells lines. (I) derived tumors were Western blotting analysis of the BACH1 protein levels in WT-HepG2, LV-Con and LV-BACH1 cell lines. (J) mages of xenograft tumors in the *HepG2* and *H-BACH1*^*−/−*^ cell lines. (K) All tumor weights for each cell group on the 27th day. (L) The growth curve of subcutaneous xenograft tumors in mice (*HepG2* and *H-BACH1*^*−/−*^ cell lines). (M) The HE-stained images of the tumors. The protein levels of KI67 in HepG2 and *H-BACH1*^*−/−*^ detected by IHC. Scale bar = 500 µm in ×40 photographs, and 100 µm in ×200 photographs. Data are shown as mean ± SD (n = 3 × 3, ^**^p < 0.01, ^***^p < 0.01, and ^****^p < 0.01).

Next, the wild type HepG2 and *H-BACH1^*−/−*^* cell lines were injected subcutaneously into nude mice to investigate the effect of BACH1 on the growth of HCC cells in vivo. Cell proliferation was assessed in vivo by measuring weights and volumes of the tumors. The size of all tumors was measured every three days until the 27nd day, when all mice were executed and the transplanted tumors were removed and weighed. Relevant results revealed that knockout of BACH1 inhibited xenograft tumors growth in mice (Figure 1J). As shown in Figure 1(K & L), the average weight and volume of tumors yielded by *H-BACH1^*−/−*^* cells were significantly lower than those of the control group on the 27th day. Moreover, histological examination unraveled that there were necrotic areas in tumor tissues derived from *H-BACH1*^*−/−*^ cells, while wild type cells did not. The expression of KI67, a cell proliferation marker was significantly decreased in *H-BACH1*^*−/−*^ derived tumors (Figure 1M). Taken together, these results suggested that knockout of BACH1 suppressed the proliferation and malignant growth of HCC cells in vivo.

### 3.2 BACH1 promoted HCC cell proliferation in vitro

The above-mentioned data indicated that knockout of BACH1 suppressed the proliferation of HCC cells in vivo. Here, we explored the biological role of BACH1 in proliferation of HCC cells in vitro. Cellular growth curves by CCK8 revealed that the proliferation capacity of *H-BACH1*^*−/−*^ and *LM3-BACH1*^*−/−*^cells was significantly reduced compared with wild-type HepG2 and HCCLM3 cells (Figure 2, A & B). However, LV-BACH1 cell proliferation capacity was significantly enhanced compared with LV-Con by ectopically expressing BACH1 (Figure 2C). The results of colony formation assays showed that knockout of BACH1 visibly inhibited clone formation rate in HepG2 cells. In contrast, the ectopic expression of BACH1 remarkably increased the rate of colony formation in LV-BACH1 cells (Figure 2, D & E). Next, we detected cell cycle changes in several different HCC cell lines by flow cytometry. The results showed that compared with wild type controls, the G0/G1 phases of both *H-BACH1*^*−/−*^ and *LM3-BACH1*^*−/−*^ cells were significantly prolonged and the G2/M phases were reduced (Figure 2F to I). It was suggested that knockout of BACH1 could arrest cell cycle at G0/G1 phase, thereby slowing down cell growth. Nevertheless, compared with LV-Con cells, the LV-BACH1 cells had a shortened S phase and a significantly longer G2/M phase (Figure 2, J & K), suggesting an increased number of cells during division phase and accelerated cell growth. Furthermore, we analyzed the effect of BACH1 on the protein expression of several genes related to tumor cell growth using western blotting. As shown in Figure 2L, knockout of BACH1 in HepG2 cells remarkably up-regulated the protein level of tumor suppressor gene p53, down-regulated epidermal growth factor receptor *LM3-BACH1*^*−/−*^ (EGFR) and vascular endothelial growth factor A (VEGFA) protein levels. In *LM3-BACH1^−/−^* cells, the EGFR protein expression level was remarkably decreased while VEGFA was only slightly down-regulated compared to control group (Figure 2M). In contrast, ectopic expression of BACH1 obviously suppressed p53 protein abundance and increased the protein abundance of cyclin A2 and EGFR in LV-BACH1 cells (Figure 2N). Altogether, these findings indicated that BACH1 facilitated the proliferation of HCC cell in vitro, knockout of BACH1 could inhibit the HCC cells growth and proliferation in vitro.

**Fig. 2.**
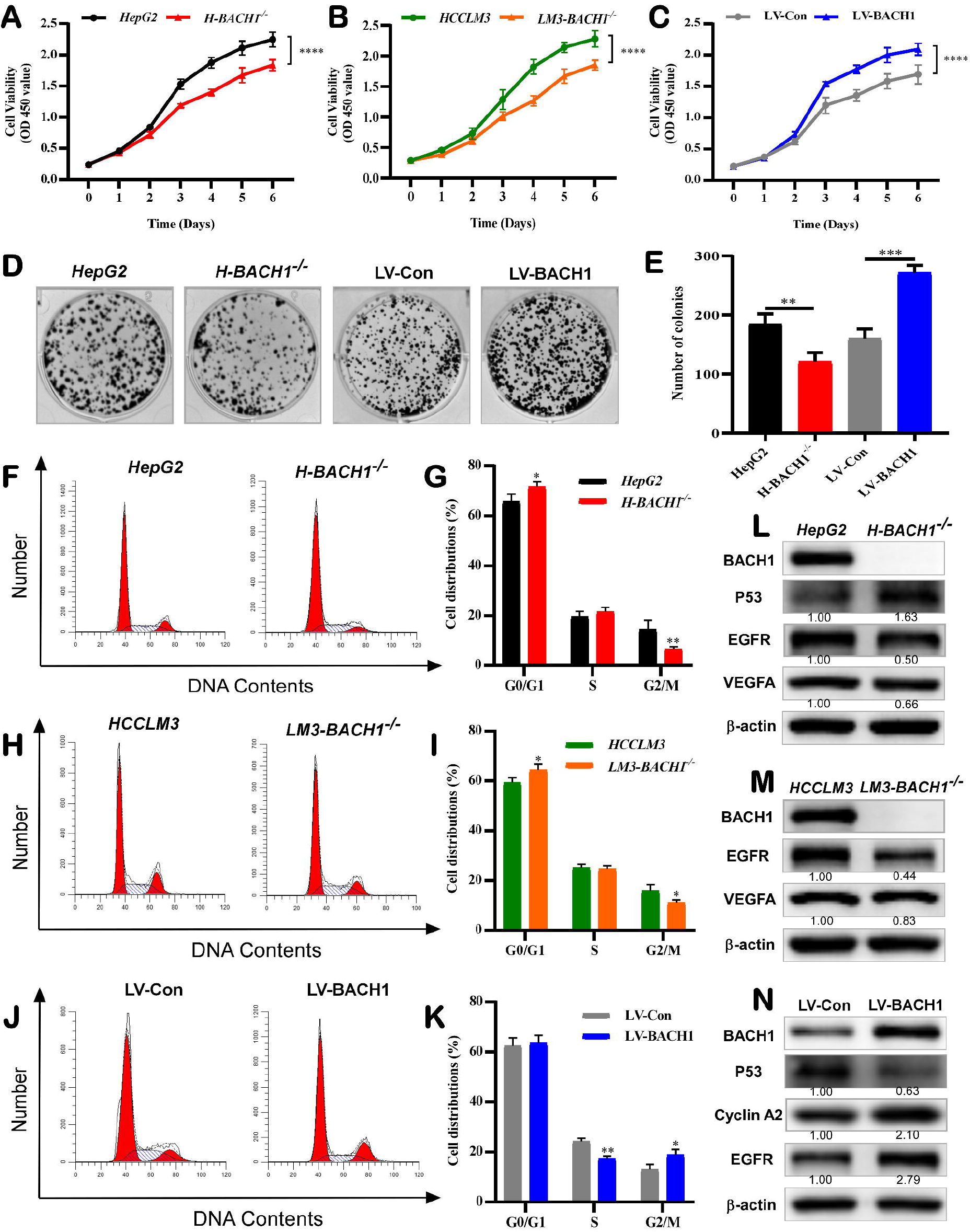
BACH1 promoted HCC cell proliferation in vitro. (A) The cellular growth curves of *HepG2* and *H-BACH1*^*−/−*^ cell lines. (B) The cellular growth curves of *HCCLM3* and *LM3-BACH1*^*−/−*^ cell lines. (C) The cellular growth curves of LV-Con and LV-BACH1 cell lines. (D, E) Colony formation in *HepG2, H-BACH1*^*−/−*^, LV-Con and LV-BACH1 cell lines, and their clone clusters were counted. (F, G) Cell cycles of *HepG2* and *H-BACH1*^*−/−*^ cell lines. (H, I) Cell cycles of *HCCLM3* and *LM3-BACH1*^*−/−*^ cell lines. (J, K) Cell cycles of LV-Con and LV-BACH1 cell lines. (L) The protein abundances of BACH1, p53, EGFR and VEGFA in *HepG2* and *H-BACH1*^*−/−*^ cell lines. (M) The protein abundances of BACH1, EGFR and VEGFA in HCCLM3 and *LM3-BACH1*^*−/−*^ cell lines. (N) The protein abundances of BACH1, p53, cyclin A2 and EGFR in LV-Con and LV-BACH1 cell lines. Data are reported as mean ± SD (n = 3 × 3, ^*^p < 0.05, ^**^p < 0.01, ^***^p < 0.01, and ^****^p < 0.01).

### 3.3 Knockout of BACH1 induced HCC cell apoptosis and ROS generation, decreased MMP

We analyzed the apoptosis of several HCC cells by flow cytometry, and results demonstrated that compared to the controls, The ratio of apoptotic cells was markedly elevated in *H-BACH1*^*−/−*^and *LM3-BACH1*^*−/−*^cells (Figure 3A to D) and obviously decreased in LV-BACH1 cells (Figure 3, E & F). Further qPCR and western blot analysis showed that knockout of BACH1 in HepG2 and HCCLM3 cell lines led to a significant increase in cysteinyl aspartate specific proteinase-3 (Caspase-3) mRNA level and the cleaved Caspase-3 protein level, a decrease in the mRNA and protein expression levels of antiapoptotic BCL2. However, no significant changes were observed in the protein and mRNA levels of proapoptotic BAX compared with control values in WT cells (Figure 3G to J). By contrast, ectopic expression of BACH1 obviously suppressed Caspase-3 mRNA abundance and cleaved Caspase-3 protein abundance, increased the mRNA and protein levels of BCL2 in LV-BACH1 cells with no significant changes in protein and mRNA abundances of BAX (Figure 3, K & L). As shown in Figure 3(A & C), knockout of BACH1 mainly promoted early apoptosis in HCC cells, and cells undergoing early apoptosis were usually accompanied by a decrease in MMP. Therefore, the changes of MMP in HepG2 and HCCLM3 cells after BACH1 knockout were detected by JC-1 staining. As expected, results showed that knockout of BACH1 reduced the MMP levels, as evidenced by the decreases of red fluorescence of JC-1 polymer in mitochondrial matrix of *H-BACH1*^*−/−*^ and *LM3-BACH1*^*−/−*^ cells, the increases of green fluorescence of JC-1 monomer compared to wild-type HepG2 and HCCLM3 cells (Figure 3, M & N). To further investigate whether knockout of BACH1-induced apoptosis was associated with intracellular ROS, we assayed ROS levels in several HCC cell lines using fluorescent probe DCFH-DA. It is shown in Figure 3 (O & P), the ROS levels of *LM3-BACH1*^*−/−*^ cells were remarkably higher than those of wild-type HCCLM3 cells, whereas compared with HepG2 cells,*H-BACH1*^*−/−*^cells had only slightly up-regulated ROS levels. In contrast, intracellular ROS of LV-BACH1 were obviously reduced compared to the LV-Con cells by ectopically expressing BACH1 (Figure 3Q). Collectively, these suggested that the induction of HCC cell apoptosis by BACH1 knockout might via mitochondrial pathway, thus suppressing tumor progression.

**Fig. 3.**
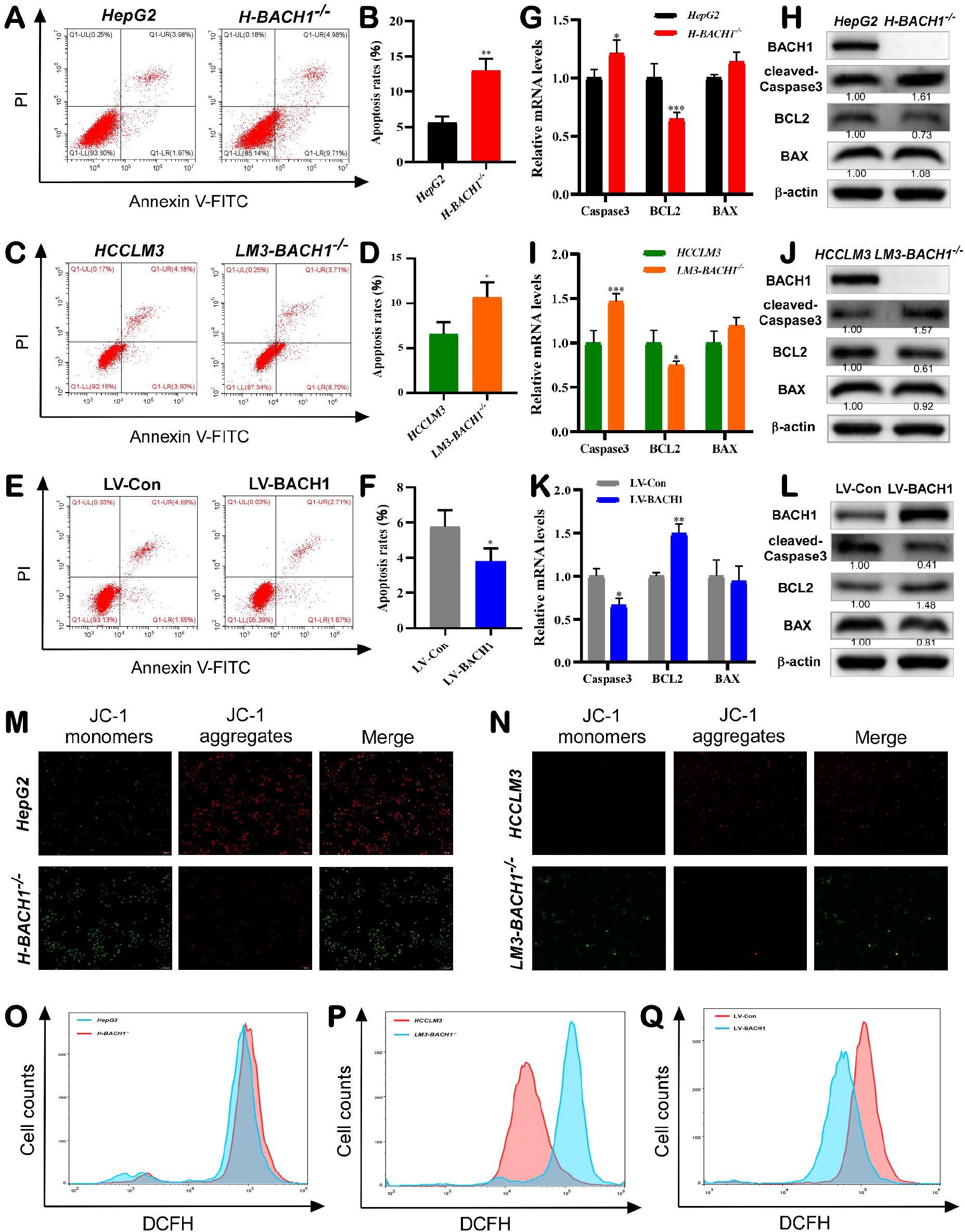
Knockout of BACH1 induced HCC cell apoptosis and ROS generation, decreased MMP. (A, B) Cell apoptosis of *HepG2* and *H-BACH1*^*−/−*^ cell lines were analyzed with flow cytometry. (C, D) Cell apoptosis of *HCCLM3* and *LM3-BACH1*^*−/−*^ cell lines were analyzed with flow cytometry. (E, F) Cell apoptosis of LV-Con and LV-BACH1 cell lines were analyzed with flow cytometry. (G) The mRNA levels of BCL2, BAX, and Caspase-3 in *HepG2* and *H-BACH1*^*−/−*^ cells. (H) After knocking out BACH1 in HepG2 cells, the changes of BCL2, cleaved Caspase-3 and BAX protein abundances. (I) The mRNA levels of BCL2, BAX, and Caspase-3 in *HCCLM3* and *LM3-BACH1*^*−/−*^ cells. (J) After knocking out BACH1 in HCCLM3 cells, the changes of BCL2, cleaved Caspase-3 and BAX protein abundances. (K) RT-qPCR analysis of Caspase-3, BCL2 and BAX mRNA levels in LV-Con and LV-BACH1 cell lines. (L) The protein *LM3-BACH1*^*−/−*^ abundances of BAX, BCL2 and cleaved Caspase-3 in LV-Con and LV-BACH1 cell lines. (M) The MMP of *HepG2* and *H-BACH1^*−/−*^* cells were measured by JC-1 staining and images were acquired using fluorescence microscopy. (N) The MMP of *HCCLM3* and *H-BACH1*^*−/−*^ cells were measured by JC-1 staining and images were acquired using fluorescence microscopy. (O) Intracellular ROS levels in *HepG2* and *LM3-BACH1*^*−/−*^ cells were assayed using fluorescent probe DCFH-DA. (P) Intracellular ROS levels in *HCCLM3* and *LM3-BACH1*^*−/−*^ and cells were assayed using fluorescent probe DCFH-DA. (Q) The ROS levels of LV-Con and LV-BACH1 cells were assayed using fluorescent probe DCFH-DA. Data are presented as mean ± SD (n = 3 × 3, ^*^p < 0.05, ^**^p < 0.01, and ^***^p < 0.01).

### 3.4 Knockout of BACH1 suppressed PI3K-AKT-mTOR and TGF-β1/SMAD signaling pathway

To gain deeper understanding of the biological function of BACH1 in HCC and its regulatory role in HCC progression, we further performed transcriptome sequencing of wild type HepG2, *H-BACH1*^*−/−*^, LV-Con and LV-BACH1 cell lines. As shown in Figure S1(A & B), principal component analysis (PCA) proved that the clear different transcriptional patterns existed between *H-BACH1*^*−/−*^ (or LV-BACH1) and HepG2 (or LV-Con) groups. The volcano map indicated that compared to HepG2, 556 DEGs were up-regulated and 482 DEGs were down-regulated in the *H-BACH1*^*−/−*^group (Figure 4A), while 540 genes were significantly up-regulated and 91 genes were significantly down-regulated in LV-BACH1 compared with the LV-Con group (Figure S1C). Next, GO analysis suggested that the top 20 enriched terms for *H-BACH1*^*−/−*^ versus HepG2 cells associated with cellular response to external stimulus, response to unfolded protein, epithelial cell development and regulation of cell growth (Figure 4B). In contrast, the top 20 enriched terms of LV-BACH1 versus LV-Con cells related to regulation of binding, reactive oxygen species metabolic process and regulation of mitochondrial membrane potential (Figure S1D). KEGG analysis showed that small cell lung cancer, PI3K-AKT signaling pathway, apoptosis, p53 signaling pathway, and MAPK signaling pathway were corporately enriched in *H-BACH1*^*−/−*^ versus HepG2 and LV-BACH1 versus LV-Con cells (Figures 4C and S1E). Notably, hepatocellular carcinoma was markedly enriched in *H-BACH1*^*−/−*^versus HepG2 cells, suggesting that BACH1 knockout might play a critical role in the HCC development and progression. Besides, further gene set enrichment analysis (GSEA) -GO revealed that multiple metabolic processes such as unsaturated fatty acid, icosanoid were activated while DNA-binding transcription activator activity was suppressed in *H-BACH1*^*−/−*^ cells (Figure 4D). By contrast, the top activated terms of LV-BACH1 versus LV-Con cells involved hormone metabolic process, while the top suppressed terms were mainly about ribosome (Figure S1F). According to the GSEA-KEGG analysis, DEGs between *H-BACH1*^*−/−*^ and HepG2 groups were enriched in five activated pathways of phagosome, linoleic acid metabolism, chemical carcinogenesis-DNA adducts, drug metabolism-cytochrome P450 and arachidonic acid metabolism (Figure 4E). In LV-BACH1 cells, cell adhesion molecules and phagosome were activated, spliceosome and ribosome pathway were suppressed (Figure S1G).

**Fig. 4.**
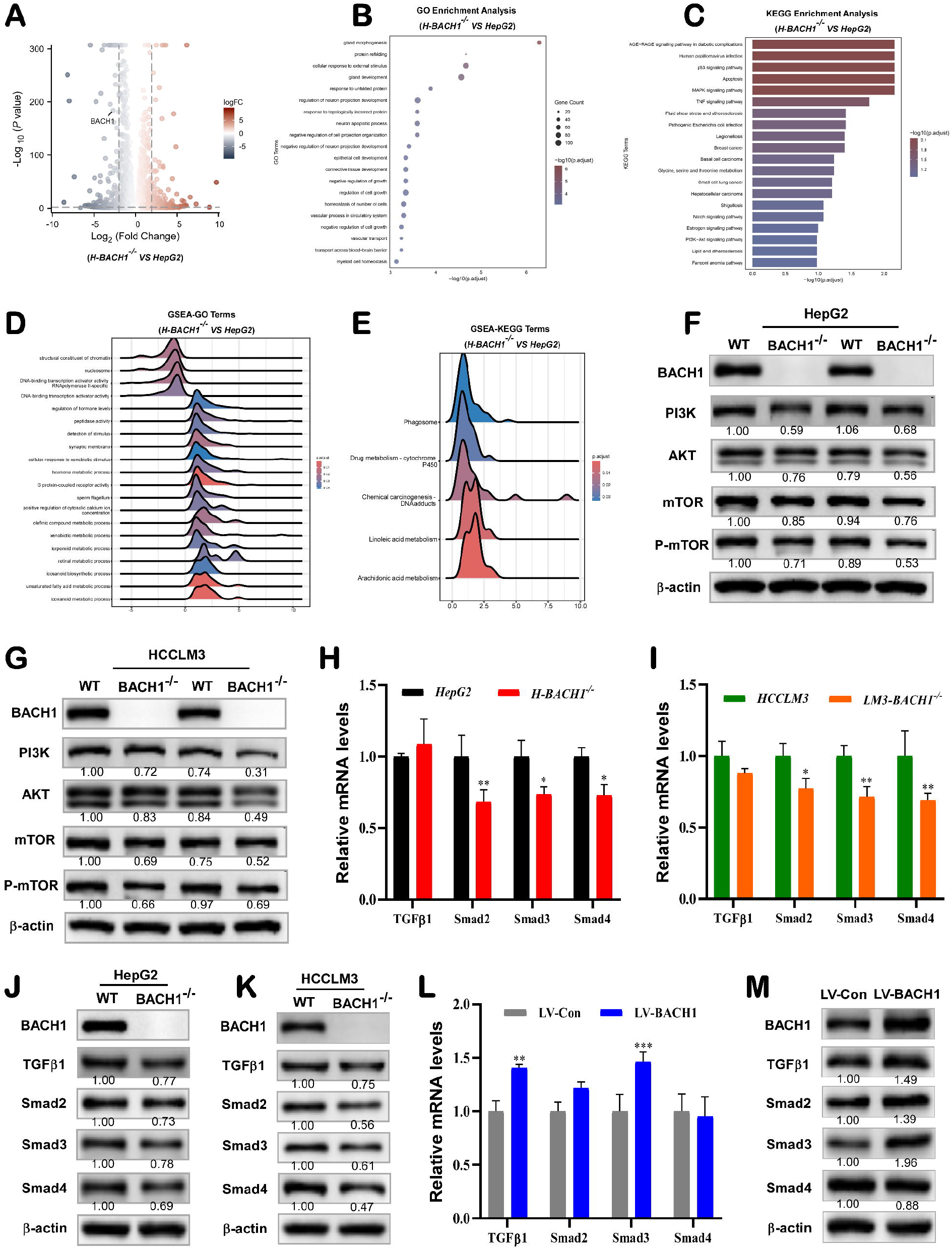
Knockout of BACH1 suppressed PI3K-AKT-mTOR and TGF-β1/SMAD signaling pathway. (A) The gradient volcano map of DEGs in *H-BACH1*^−/−^ versus HepG2 cell lines. (B) GO enrichment analysis of DEGs in *H-BACH1*^−/−^ versus HepG2 cell lines. (C) KEGG enrichment analysis of DEGs between *H-BACH1*^−/−^ and HepG2 cell lines. (D) The ridge plot of activated and suppressed terms based on GSEA-GO analysis in *H-BACH1*^−/−^ versus HepG2 cell lines.(E) The ridge plot of activated five pathways based on GSEA-KEGG analysis in *H-BACH1*^−/−^ versus HepG2 cell lines. (F) Western blotting analysis of the BACH1, AKT, PI3K, mTOR, and p-mTOR protein levels in *HepG2* and *H-BACH1*^−/−^ cells. (G) Western blotting analysis of the BACH1, AKT, PI3K, mTOR, and p-mTOR protein levels in *HCCLM3* and *LM3-BACH1*^−/−^ cells. (H) The mRNA levels of TGF-β1, SMAD2, SMAD3 and SMAD4 in *HepG2 H-BACH1*^−/−^ cells. (I) The mRNA levels of TGF-β1, SMAD2, SMAD3 and SMAD4 in *HCCLM3* and *LM3-BACH1*^−/−^ cells. (J) The effects of BACH1 knockout on TGF-β1, SMAD2, SMAD3 and SMAD4 protein abundances in HepG2 cell line. (K) The effects of BACH1 knockout on TGF-β1, SMAD2, SMAD3 and SMAD4 protein abundances were examined by Western blotting in HCCLM3 cell line. (L) RT-qPCR analysis of TGF-β1, SMAD2, SMAD4 and SMAD3 mRNA levels in LV-Con and LV-BACH1 cells. (M) The protein levels of BACH1, TGF-β1, SMAD2, SMAD3 and SMAD4 in LV-Con and LV-BACH1 cells. Data are reported as mean ± SD (n = 3 × 3, ^*^p < 0.05,^**^p < 0.01, and ^***^p < 0.01).

Based on the results of transcriptome sequencing analysis, Western blotting was used to validate the regulation of BACH1 on PI3K-AKT pathway. In Figure 4 (F & G), the protein abundances of PI3K and AKT were decreased in both *H-BACH1*^*−/−*^and *LM3-BACH1*^*−/−*^cell lines compared with controls. Knockout of BACH1 significantly down-regulated the phosphorylation of mTOR protein in *H-BACH1*^*−/−*^ cells and mTOR and p-mTOR protein levels in *LM3-BACH1*^*−/−*^ cells. Also, we analyzed the effect of BACH1 on cell growth related TGF-β1/SMAD signaling pathway. The results of qPCR and western blotting demonstrated that BACH1 knockout in two HCC cell lines led to obvious downregulation on mRNA and protein levels of SMAD2, SMAD3 and SMAD4. Meanwhile, knocking out BACH1 also reduced the protein level of TGF-β1, but did not influence its mRNA level (Figure 4H to K). In contrast, the ectopic expression of BACH1 markedly increased TGF-β1 and SMAD3 mRNA abundances, while no significant alterations were observed in the mRNA levels of SMAD2 and SMAD4 (Figure 4L). Compare with LV-Con cells, protein levels of TGF-β1, SMAD2 and SMAD3 were observably up-regulated in LV-BACH1 cells, while no significant changes in the SMAD4 protein abundance (Figure 4M).

### 3.5 BACH1 accelerated cell proliferation by targeting the PDP1-PI3K/AKT/mTOR axis

Tumor cells autonomously change metabolic fluxes according to various pathways to address the energetic and material requirements for rapid proliferation[22, 23]. PDP1 catalyzes dephosphorylation to activate PDC, promoting carbohydrate utilization and cellular energy production. Our study revealed that the protein and mRNA levels of PDP1 were obviously down-regulated in *H-BACH1*^*−/−*^ and *LM3-BACH1*^*−/−*^cells (Figure 5A to D). In stark contrast, PDP1 protein and mRNA levels in LV-BACH1 cells were significantly elevated by ectopic expression of BACH1 compared with the control (Figure 5, E & F). To further explore the regulatory of BACH1 on PDP1 transcription, promoter region for PDP1 gene was embedded in the pGL3-Basic vector (Figure 5G), and performed dual luciferase reporter gene analysis. Results indicated that BACH1 remarkably enhanced the transcriptional activity of PDP1 promoter in both HEK293 and HepG2 cells (Figure 5, H & I). Next, sequence analysis revealed the presence of several ARE consensus DNA-binding sites in PDP1 promoter region (Figure 5J). Distinct ARE-driven luciferase reporter genes were trans-activated by BACH1 in HEK293 or HepG2 cells (Figure 5, K & L), suggesting that BACH1 bound to the ARE sites in promoter region of PDP1 to activate its transcriptional expression. Besides, three different siRNA pairs were synthesized for silencing PDP1 gene expression to delve its role in BACH1 regulation of HCC progression. As shown in Figure 5(M & N), the mRNA expressions of PDP1 in HepG2 and HCCLM3 cells were markedly reduced by all three siPDP1s, while its protein level was not remarkably downregulated by siPDP1-3 in HepG2 cells (Figure 5O). Such being the case, we selected two siRNAs (siPDP1-1 and siPDP1-2) that reduced the protein levels of PDP1 in both two HCC cells for subsequent experiments (Figure 5, O & P). Western blotting results indicated that the protein abundances of AKT, PI3K, mTOR and phosphorylation of mTOR protein were remarkably downregulated by silencing PDP1 in HepG2 cells (Figure 5Q). In HCCLM3 cells, knockdown of PDP1 resulted in a significant decrease in PI3K, AKT protein levels and phosphorylation of mTOR protein, but did not affect mTOR protein level (Figure 5R). Of note, PDP1 functioned as a tumor promoter, because the clonogenicity of HepG2 cells was suppressed by reducing its expression levels (Figure 5S). Altogether, BACH1 targeted PDP1-PI3K/AKT/mTOR axis to accelerate HCC cell proliferation.

**Fig. 5.**
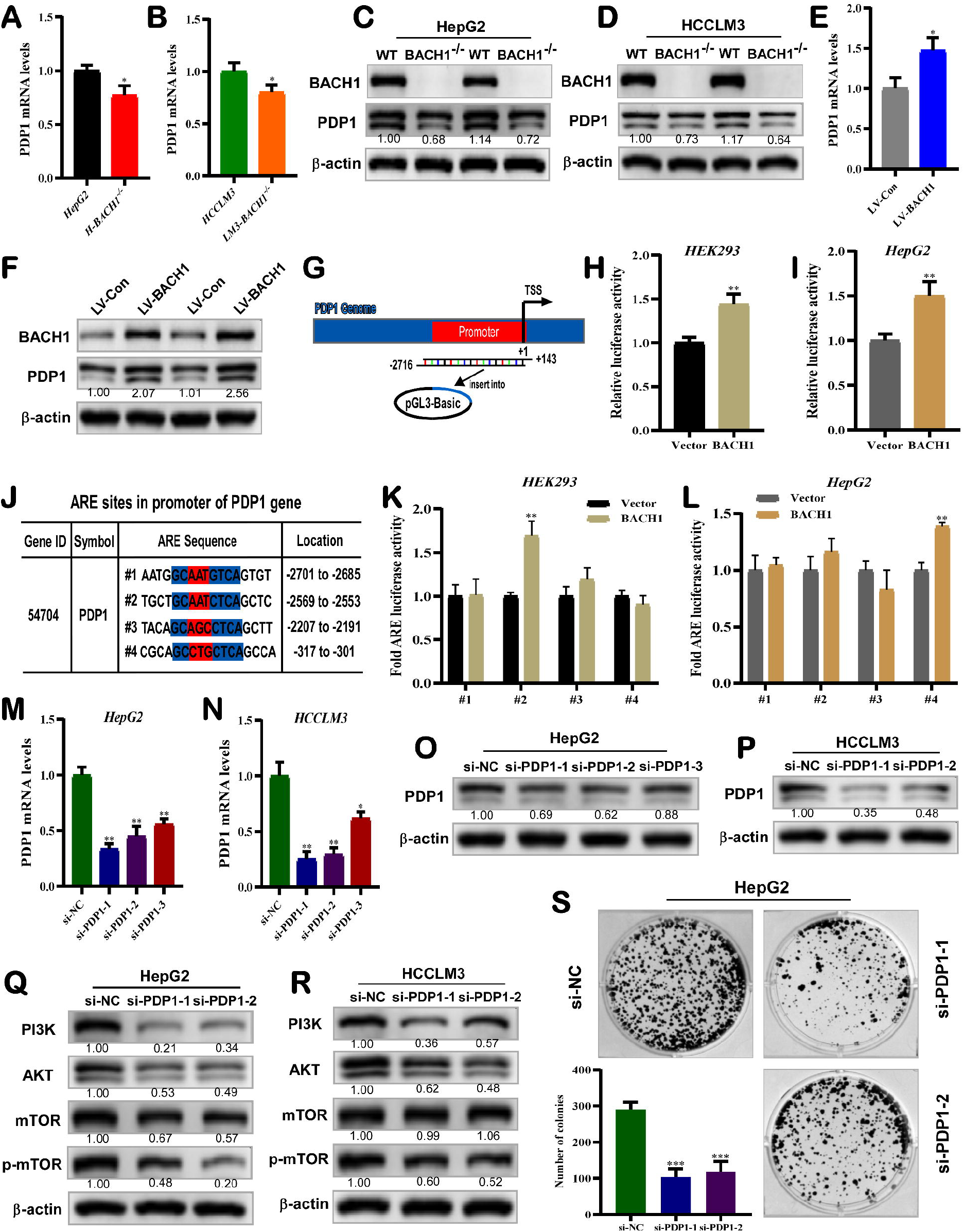
BACH1 accelerated cell proliferation by targeting the PDP1-PI3K/AKT/mTOR axis. (A) The mRNA levels of PDP1 in *HepG2* and *H-BACH1*^−/−^ cells. (B) The mRNA levels of PDP1 in *HCCLM3* and *LM3-BACH1*^−/−^ cells. (C) The PDP1 protein abundances in *HepG2* and *H-BACH1*^−/−^ cells. (D) The PDP1 protein abundances in *HCCLM3* and *LM3-BACH1*^−/−^ cells. (E) The PDP1 mRNA levels in LV-Con and LV-BACH1 cells. (F) The protein levels of PDP1 in LV-Con and LV-BACH1 cells. (G) Diagram of the PDP1 promoter reporter gene plasmid. Its promoter region was also indicated. (H) The influence of BACH1 on transcriptional activation of PDP1 promoter by Luciferase reporter gene assay in HEK293 cells. (I) The effect of BACH1 on transcriptional activity of PDP1 promoter was detected by Luciferase reporter gene assays in HepG2 cells. (J) The ARE consensus DNA-binding sites in the promoter of PDP1 were listed herein. (K) The effect of BACH1 on distinct ARE luciferase activities in PDP1 promoter were detected in HEK293 cells. (L) The effect of BACH1 on distinct ARE luciferase activities in PDP1 promoter were detected in HepG2 cells. (M) After transfection of three PDP1-targeting siRNAs (siPDP1-1, siPDP1-2 and siPDP1-3) in HepG2 cells, the PDP1 mRNA levels were measured. (N) After transfection of three PDP1-targeting siRNAs (siPDP1-1, siPDP1-2 and siPDP1-3) in HCCLM3 cells, the PDP1 mRNA levels were measured. (O) After transfection of siPDP1-1, siPDP1-2 and siPDP1-3 in HepG2 cells, the abundance of PDP1 protein was determined. (P) After transfection of siPDP1-1 and siPDP1-2 in HCCLM3 cells, the abundance of PDP1 protein was determined. (Q) HepG2 cells were transfected by siPDP1-1 and siPDP1-2, then PI3K, AKT, mTOR and p-mTOR protein levels were measured. (R) HCCLM3 cells were transfected by siPDP1-1 and siPDP1-2, then PI3K, AKT, mTOR and p-mTOR protein levels were measured. (S) Colony formation in HepG2 cells transfected with si-NC, siPDP1-1 and siPDP1-2. Their clone clusters were enumerated. All data are shown as mean ± SD (n = 3 × 3, ^*^p < 0.05, ^**^p < 0.01, and ^***^p < 0.01).

## 4. Discussion

During several stages of cancer progression, cancer cells need to continuously express or activate transcription factor to accelerate growth and survivability [24]. In recent years, the function of CNC-bZIP transcription factor BACH1, in multiple malignancies has been adequately characterized. Previous studies showed that BACH1 expression was up-regulated within HCC sample, its higher levels were correlated with poorer overall patient survivals. Furthermore, it accelerated HCC cell invasion and metastasis through activating motility-associated PTK2 as well as IGF1R gene [11]. Here, our study verified functions of BACH1 in regulating HCC progression, and further explored the underlying molecular mechanisms. The results indicated that knockout of BACH1 significantly diminished malignant proliferation of HCC cells, whereas overexpression of BACH1 facilitated HCC cell growth. Mechanistically, knockout of BACH1 induced cell apoptosis via mitochondrial pathway and suppressed cell growth related TGF-β1/SMAD signaling pathway, thereby suppressing tumor progression. On the other hand, BACH1 accelerated HCC cell proliferation by targeting the PDP1-PI3K/AKT/mTOR axis (Figure 6).

**Fig. 6.**
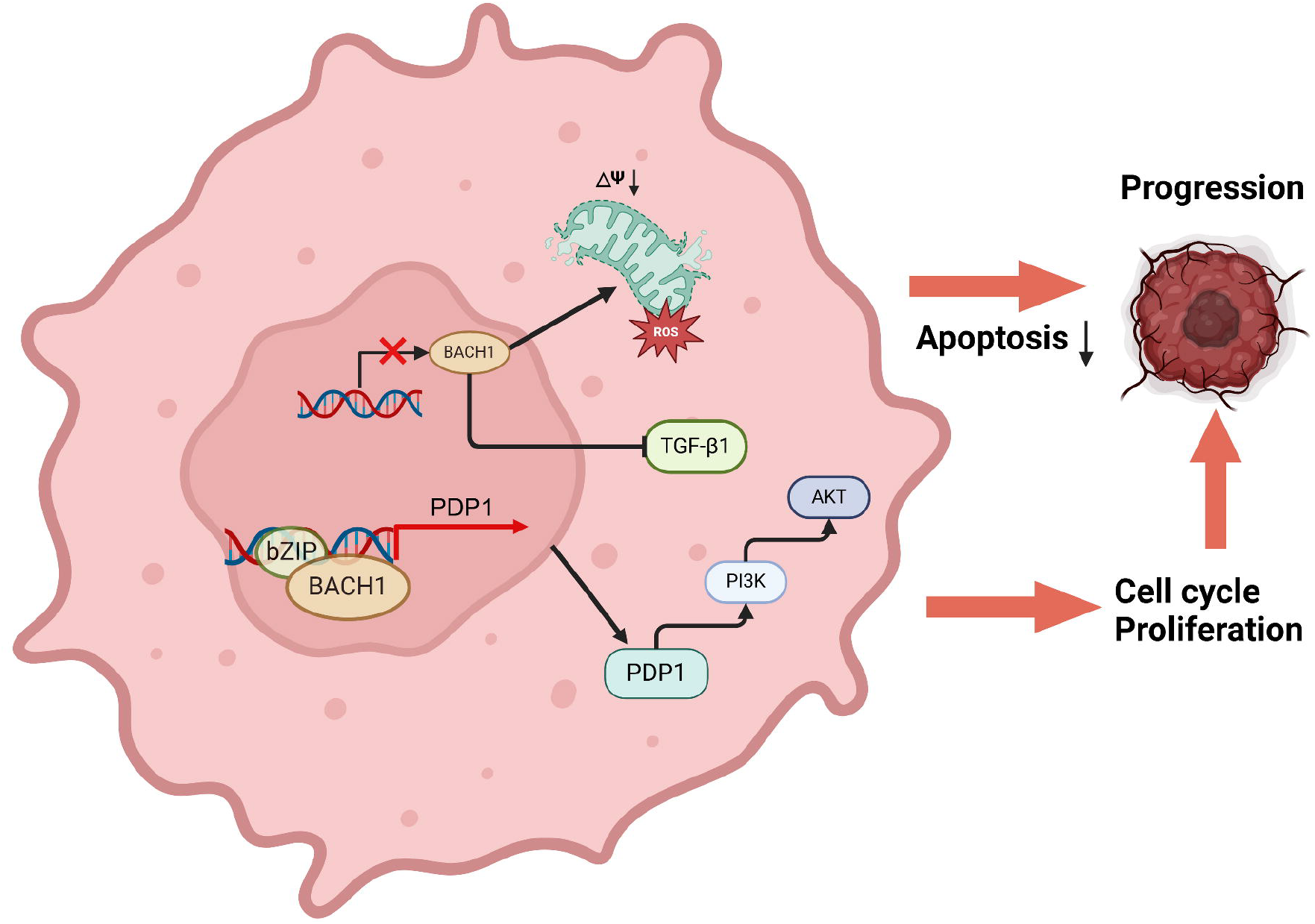
Schematic presentation of mechanisms that BACH1 promotes the progression of HCC. BACH1 activates transcription of PDP1, and accelerates HCC cell proliferation by targeting the PDP1-PI3K/AKT/mTOR axis. BACH1 also facilitates the progression of cell cycle and suppresses apoptosis, thus facilitating HCC progression. Knockout of BACH1 induced HCC cell apoptosis via mitochondrial pathway and suppressed cell growth related TGF-beta1 pathway, thus suppressing tumor progression.

In detail, our results revealed that knockout of BACH1 dramatically suppressed the growth of xenograft neoplasms in nude mouse, which is consistent with previous studies of BACH1 knockdown in HCC cells [11]. Vitro assays demonstrated that HCC cell proliferation, clonogenicity and cell cycle turnover were promoted by BACH1 overexpression, but the opposite effects were manifested by BACH1 knockout. Of note, BACH1 prominently downregulated the protein expression of p53 gene, while upregulated EGFR. p53 is known to be an essential tumour suppressive factor which mutates across over one-half of human cancers [25]. KEGG analysis indicated that p53 signaling pathway was significantly enriched upon both knockout and overexpression of BACH1 in HCC cells. Previous studies have found BACH1 negatively regulated p53 pathway, that was consistent with our results [26]. Moreover, EGFR is a receptor tyrosine kinase whose abnormal expression has been linked to a variety of cancers [27]. The EGFR pathways have been demonstrated involvement in HCC malignant progression [28, 29], suggesting BACH1 may promote HCC progression by mediating p53 or EGFR signaling pathways. The above findings prove that BACH1 acts as a dominant tumour-promoting factor which has vital functions in the HCC progression.

Apoptosis suppresses tumorigenesis and progression at multiple stages, with the mitochondrial pathway is the commonest form for cell death in cancer [30]. BACH1 was found to inhibit apoptosis induced by lncRNA AC016727.1 knockdown of human lung cancer cells [31]. Furthermore, loss of BACH1 in both mantle cell lymphoma and lung cancer stem cells markedly triggered cell apoptosis [32, 33]. Nonetheless, the potential roles for BACH1 in HCC cell apoptosis remain largely unknown. Our results demonstrated that knockout of BACH1 observably induced HCC cell apoptosis, while overexpression of BACH1 had the opposite effect. Mitochondria plays a central role in intrinsic apoptosis. The pathway is principally regulated by BCL2 family of proteins and a high BAX/BCL2 expression ratio stimulates apoptosis through increasing mitochondrial membrane permeability [34, 35]. Our study found that BACH1 knockout affected mitochondrial function by down-regulating the expression of antiapoptotic BCL2 and reducing mitochondrial membrane potential, accompanied by increased ROS generation. Meanwhile, Caspase3 was activated after BACH1 knockout, ultimately triggering apoptosis. All these results suggested that knockout of BACH1 induced HCC cell apoptosis via mitochondrial pathway.

Dysregulated PI3K-AKT-mTOR pathways are very frequent in HCC, this signaling network controls a majority of cancer features which include cell cycle, proliferation, metastasis and metabolism etc. [36]. Based on transcriptome sequencing, we further verified the regulation of BACH1 on PI3K-AKT signaling pathway enriched by KEGG analysis and its mechanistic target of rapamycin (mTOR). The results showed that knockout of BACH1 could suppress PI3K-AKT-mTOR signaling. BACH1 also mediated the regulation of TXN on HCC stemness through activating AKT-mTOR pathway [16], suggesting that BACH1 promotes PI3K-AKT-mTOR signalling to accelerate HCC cell self-renewal and malignant growth. Besides, TGF-β/SMAD pathway also plays critical roles in facilitating growth and motility. TGF-β1 was found to stimulate proliferation of hepatoma cells by increasing the expression of glioma-associated oncogene homolog 1 [37, 38]. Our studies indicated that overexpression of BACH1 significantly activated the TGF-β1/SMAD signaling pathway, whereas knockout of BACH1 suppressed TGF-β1/SMAD signaling. Collectively, BACH1 enhances PI3K-AKT-mTOR and TGF-β1/SMAD signaling, promotes the progression of HCC.

PDP1 is an important phosphatase known for its involvement in cellular energy metabolism. Previous studies have showed that PDP1 activates PDC to accelerate tumor growth in human pancreatic adenocarcinoma and prostate cancer [18, 20], but impede lung cancer [39]. Nevertheless, the function of PDP1 in HCC is poorly described. Our results demonstrated PDP1 manifested as a tumor promoter, because the clonogenicity of hepatoma cells was suppressed by reducing its expression level. BACH1 bound to the ARE sites in promoter region of PDP1, so to activate its transcription. Interestingly, like BACH1, down-regulation of PDP1 in HCC cells also suppressed the PI3K-AKT-mTOR signaling. In pancreatic adenocarcinoma, mTOR activation contributed to PDP1-driven proliferation of tumour cells [18]. These suggest that PDP1 mediates the regulation of BACH1 on HCC progression. However, some limitations of this study should be pointed out. The role of PDP1 as a key enzyme regulating PDC activity in HCC metabolism has not been clarified. Moreover, reduction of PDC activity is considered to in part promote the Warburg effect, and its tumor-promoting or suppressive effect in cancer remains controversial [20, 40-42]. In further studies, we will focus on whether PDP1 also contributes to Bach1-driven malignant proliferation of HCC cells in an enzymatic-dependent manner.

In summary, our study highlights that BACH1 manifests as a robust tumor promotor which promotes HCC progression via targeting PDP1-PI3K/AKT/mTOR signaling pathway. This provides potential strategies for developing new clinical options for the treatment of HCC.

## Author Contributions

Q.B. wrote the original draft and devised most of experiments. X.Q. and Q.W. contributed to the data analysis and animal experiments. S.L. provided valuable advices for this work. S.L. and Z.W. collected tissue samples and performed histological examinations. Finally, Y.Z. guided this work and modified the present manuscript. All authors read and approved the final version of the manuscript to be published.

## Ethics statement

The animal experiments were approved by Chongqing University Laboratory Animal Welfare and Ethics Committee.

