## Supplemental for "BACH1 promotes hepatocellular carcinoma progression by targeting PDP1 towards the PI3K-AKT-mTOR signaling activation"

**Supplementary materials and methods**

**Cell line culture and transfection**

The human hepatocellular carcinoma (HepG2) cell line was obtained originally from ATCC (Zhong Yuan Ltd., Beijing, China). Additional three cell lines, respectively *H-BACH1^−/−^* (with a knockout mutant), Lentiv-PLJM (control) or Lentiv-BACH1 (stably overexpressing BACH1) were created from HepG2 cells. In addition, HCCLM3 cell line was obtained from the Live Cancer Institute (Fudan University of China) and another, *LM3-BACH1^−/−^* (with a knockout mutant) was created from it.

All experimental cell lines were cultivated in DMEM (GIBCO, Life technologies) supplemented with 10% fetal bovine serum (FBS, Biological Industries, Israel), penicillin and streptomycin (100 units/mL, Beyotime, Shanghai, China), in the 37°C incubator with 5% CO_2_. The cDNA sequences of human BACH1 gene was cloned into pcDNA3.1 vector to construct BACH1 expression plasmid. Three siRNA pairs targeting different nucleotide sequences of PDP1 (Table S1) were synthesized for silencing its gene expression. Each of these plasmid or siRNAs was transfected into cells by incubating with Lipofectamine 3000 (Invitrogen, Carlsbad, CA, USA).

**The *BACH1* gene-editing by CRISPR/Cas9 to yield** ***H-BACH1^−/−^* and** ***LM3-BACH1^−/−^***

The gRNA-target sequences were designed online (http://crispr.dbcls.jp/) (Table S2) and then cloned into the Cas9/Grna (puro-GFP) Vector (Wiewsolid Biotech, [China](javascript:;)). The indicated plasmids were confirmed by sequencing and co-transfected into HepG2 or HCCLM3 cells. Subsequently, the transfected cells were screened with puromycin (Solarbio, [Beijing](javascript:;), [China](javascript:;)), diluted and inoculated into 96-well cell culture plates (with a probability of one cell per well). The positively-selected monoclonal cell lines were subjected to the genomic DNA extraction and PCR amplification of the gRNA-target-adjoining sequences to identify their genotypes, called *H-BACH1^−/−^* and *LM3-BACH1^−/−^*, respectively.

**BACH1-overexpressing and** **control cell lines were established by lentivirus**

The cDNA sequence of human BACH1 gene was cloned into the PLJM1-EGFP vector, named pLJM1-BACH1, that was verified by sequencing. Respectively, the PLJM1-BACH1 or PLJM1-EGFP, together with the virus-packaging plasmids psPAX2 and pMD2G, were co-transfected into 293T cells for 8 h, before these cells were allowed for a 24-h recovery from transfection in a complete medium containing 10% FBS. The cells continued to culture for additional 24 h and their supernatants were collected to obtain a certain amount of lentivirus. The lentivirus titer was evaluated, prior to an efficient infection of the BACH1-expressing or vectorial lentivirus into HepG2 cells. The positive monoclonal cells (called LV-BACH1 and LV-Con) were selected and saved for subsequent experiments.

**Real-time quantitative PCR**

Total RNAs of experimental cells were extracted with the RNA simple kit (Tiangen, Beijing, China) and added the premixed reverse transcription reagent (PrimeScript^™^ RT Master Mix, RR036A, Takara, JPN) to obtain the cDNAs.

The real-time PCR reaction mixture (20μL) including GoTaq qPCR Master Mix (Promega, USA), primers (synthesized by Tsingke, Chengdu, China and listed in Table S3) and cDNA templates were incubated at 95°C for 5 min, followed by 40 cycles at 95°C for 15 s and extending at 60°C for 30 s, in the CFX Connect Real-Time PCR Detection System (Bio-Rad, CA, USA). Therein, β-actin was used as an internal control for normalization. The 2-^△△^Ct method was used to calculate the relative mRNA expression abundances.

**Western blotting with distinct antibodies**

The experimental cells were lysed in the RIPA buffer (R0010, Solarbio, [Beijing](javascript:;), [China](javascript:;)) supplemented with the protease and Protein Phosphatase inhibitor (P6730, P1260, Solarbio). The lysates were diluted with 5 × SDS-PAGE loading buffer (P1040, Solarbio), denatured at 100°C for 10 min. Equal amounts of protein extracts were loaded in each well of SDS-PAGE gels containing 8% or 10% polyacrylamide, and transferred to the polyvinylidene fluoride membranes (Millipore Co., Tullagreen, Ireland). The protein-loaded membranes were immunoblotted with each of the primary antibodies, including BACH1 (sc-271211, from Santa Cruz Biotechnology), PI3K (ab151549), mTOR (ab32028), p-mTOR (ab109268), TGF-β1 (ab179695), Smad2 (ab40855), Smad3 (ab40854), Smad4 (ab40759), VEGFA (ab214424) (all eight antibodies purchased from Abcam), AKT (#4691, from [Cell Signaling Technology](http://www.baidu.com/link?url=oa-4pfbyM1Qet1G382TDQR9VSOYgUjCY4cPu1nTH9TQvcnz9xEG5djNjbxswYCoC)), p53 (#2524, from [Cell Signaling Technology](http://www.baidu.com/link?url=oa-4pfbyM1Qet1G382TDQR9VSOYgUjCY4cPu1nTH9TQvcnz9xEG5djNjbxswYCoC)), Cyclin A2 (A19036), EGFR (A11351), Caspase3 (A16793), BCL2 (A0208), BAX (A12009) (these antibodies purchased from ABclonal, Wuhan, [China](javascript:;)), PDP1 (21176, from Proteintech Group) and β-actin (TA-09, from ZSGB-BIO, [Beijing](javascript:;), [China](javascript:;)) overnight at 4°C and then the secondary antibodies for 2 h at 37°C. The immunoblots were exposed to the ECL light system and calculated by using the ImageJ software.

**Subcutaneous** **tumor xenografts in** **nude mice**

The wild type HepG2 and its derived *H-BACH1^−/−^* [cell](javascript:;) lines in exponential growth phase (1×10^7^) were suspended in 0.1 mL PBS solution and injected subcutaneously into 5-week-old male nude mice (BALB/C^nu/nu^, from HFK Bioscience, Beijing, China) to establish mouse xenograft tumor models (n = 4 per group). The tumor sizes were measured every three days and calculated by a standard formula (i.e., V = ab^2^/2). At the end of the experiment, all mice were sacrificed and their transplanted tumors were excised and weighed. Subsequently, the tumor tissues were fixed with paraformaldehyde for hematoxylin-eosin staining and Immunohistochemistry (IHC). Notably, all mice were maintained under standard diets and living conditions. All relevant experimental protocols were approved by Chongqing University Laboratory Animal Welfare and Ethics Committee.

**Immunohistochemistry (IHC)**

In brief, the fixed tumor tissues were embedded in paraffin and made into 5 μm sections. After deparaffinized with xylene and rehydrated with graded ethanol, the sections were incubated with 3% hydrogen peroxide for 10 min to block endogenous peroxidase. Subsequently, sections were microwaved in 0.01M citrate buffer (PH 6.0) for 20 min at 95°C and blocked with BSA for 30 min after cooling. Incubations were performed with primary antibodie against ki67 (abcam, ab15580) overnight at 4°C and then HRP-conjugated secondary antibodie at room temperature for 1 h. After stained with DAB, counterstained with hematoxylin, sections were dehydrated rapidly and sealed with neutral resin. Images were captured with confocal microscopy (Olympus).

**Cell Counting Kit 8 (CCK8) and colony formation assays**

Cell proliferation was assessed using CCK-8 (BG0025, Bioground, Chongqing, China) according to the manufacturer's instructions. Experimental cells (2 × 10^3^ /well) with three duplications were seeded in 96-well plates, and incubated under aseptic conditions at 37 °C with 5% CO_2_ for 0 to 6 days, respectively. Subsequently, the absorbance was measured at 450 nm wavelength using a microplate reader (iMark, Bio-Rad).

The colony formation assay was to culture the experimental cells (1000 cells/well seeded in 6-well plates) at 37 °C with 5% CO_2_ for 14 days. The cells were subjected to fixation by 4% [paraformaldehyde](https://www.sciencedirect.com/topics/medicine-and-dentistry/paraformaldehyde) and stained with crystal violet staining solution (Beyotime, Shanghai, China). Subsequently, the colony number was counted.

**Analysis of cell cycle and apoptosis by flow cytometry**

The experimental cells were digested by trypsin and suspended in pre-cooled phosphate-buffered saline (PBS), subjected to measuring cell cycle (C1052, Beyotime, Shanghai, China) or apoptosis (C1062S, Beyotime, Shanghai, China) according to the manufacturer's instructions. Cells were fixed with 70% ethanol overnight at 4°C. Subsequently, the cells were incubated with RNase A and propidium iodide (PI)-staining solution at 37°C for 30min before analysis of the cell cycle by flow cytometer. Cells were incubated in the dark with annexin V-FITC and PI at room temperature for 15 min, cell apoptosis was measured immediately by flow cytometry.

**Mitochondrial membrane potential (MMP) measurement**

Enhanced mitochondrial membrane potential assay kit with JC-1 (C2003S, Beyotime, Shanghai, China) was used to measure MMP according to the manufacturer's instructions. In brief, experimental cells were seeded in 6-well plates and incubated with JC-1 dye for 20 min at 37°C. Subsequently, cells were washed twice with JC-1 dye buffer and images were captured using a fluorescence microscope (NIKON ECLIPSE TI-SR, Japan).

**Intracellular ROS measurement**

Intracellular ROS was measured using ROS assay kit (S0033, Beyotime, Shanghai, China). Briefly, experimental cells were collected and incubated with 10 μM DCFH-DA at 37°C for 20 min. After cells were washed three times using serum-free culture medium, the DCF fluorescence intensity was measured by a flow cytometer in 488nm excitation and 525nm emission lights.

**Analysis of** **transcriptome sequencing**

Total RNAs were extracted from *WT-HepG2, H-BACH1^−/−^*, Lentiv-PLJM and Lentiv-BACH1 [cell](javascript:;) lines using TRIzol reagent (RNAiso Plus, 9108, Takara, JPN) and transcriptome sequencing was performed by Genomics Institute (Beijing, China) on the DNBSEQ platform. After the raw data were filtered, those clean reads are obtained and mapped to the reference sequences of Homo sapiens’ genome (GCF_000001405.39_GRCh38.p13) using HISAT [22]. The gene expression levels were calculated by using the RSEM method [23], and DESeq2 tool was used to identify the differentially expressed genes (DEGs), with a criteria Log_2_ fold-changes ≥1 and Q-value ≤0.01. Those DEGs were subjected to both the Gene Ontology (GO)[24] functional enrichment analysis and the Kyoto Encyclopedia of Genes and Genomes (KEGG) [25] pathway enrichment analysis.

**Dual-luciferase reporter gene assay**

The promoter sequences for PDP1 gene were inserted into the pGL3-Basic vector to construct promoter reporter gene plasmid. The ARE and adjacent sequences contained in the PDP1 promoter were inserted into the pGL3-promoter vector to construct ARE reporter gene plasmids. The construction of BACH1 expression plasmid was described in 2.1. HEK293 or HepG2 cells were seeded in 12-well plate and cultured overnight. When the cells density reached 70%-80%, they were co-transfected with BACH1 expression plasmids, reporter gene plasmids and Renilla luciferase reporter plasmid (pRL-TK) for 8h. After being incubated in complete medium for 24h, the cells were lysed and dual-reporter assay (Promega, USA) was used to assay the luciferase activities. The ratio of Firefly/Renilla luciferase activity was calculated.

**Statistical analysis**

Statistical analyses were performed using the Origin8.0 tool. All relevant data in this study were obtained from at least three independent experiments and presented as fold changes (mean ± SD). The one-way ANOVA was used to calculate the statistic differences between the various experimental groups and within groups. A P value lower than 0.05 was considered statistically significant.

**Supplementary Figure**

**Figure S1**

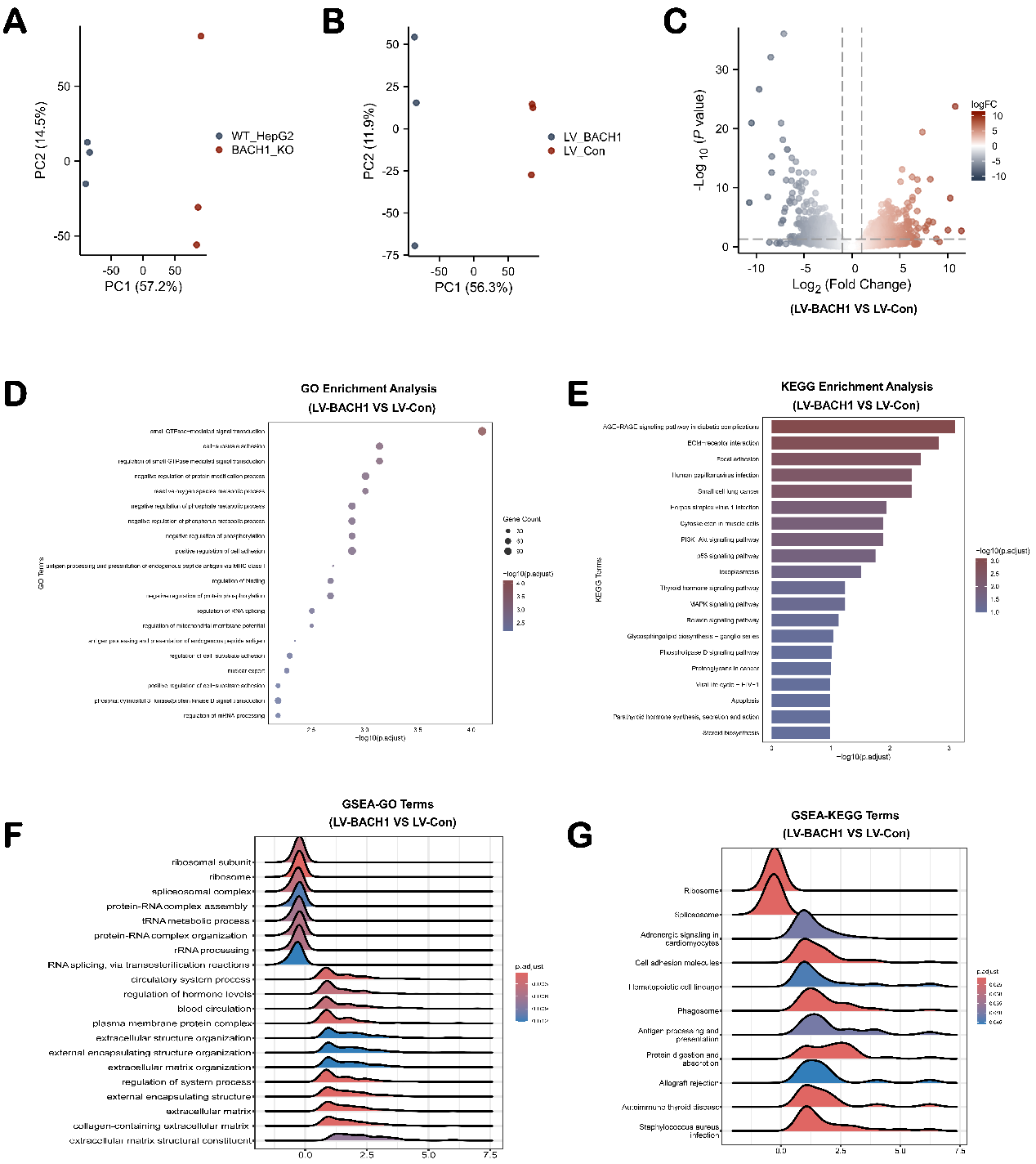

**Fig. S1. Transcriptome sequencing analysis.**

(A) The PCA analysis of HepG2 and *H-BACH1^−/−^* cell samples.

(B) The PCA analysis of LV-Con and LV-BACH1 cell samples.

(C) The gradient volcano map of DEGs in LV-BACH1 versus LV-Con cell lines.

(B) GO enrichment analysis of DEGs in LV-BACH1 versus LV-Con cell lines. The top 20 GO terms are shown in bubble chart.

(C) KEGG enrichment analysis of DEGs between LV-BACH1 and LV-Con cell lines. The bar chart shows the top 20 significantly enriched pathways.

(D) The ridge plot of activated and suppressed terms based on GSEA-GO analysis in LV-BACH1 versus LV-Con cell lines.

(E) The ridge plot of activated and suppressed pathways based on GSEA-KEGG analysis in LV-BACH1 versus LV-Con cell lines.

**Supplementary Tables**

**Table S1**. The siRNA sequences used in this study

| **siRNA** | **Nucleotide sequence (5′→ 3′)** |
| --- | --- |
| siPDP1-1 | Sense: CACCCGAUUUCCUAAUGUAdTdT |
| Antisense: UACAUUAGGAAAUCGGGUGdTdT | |
| siPDP1-2 | Sense: GGAAUCGUACUUCUUAUUUdTdT |
| Antisence: AAAUAAGAAGUACGAUUCCdTdT | |
| siPDP1-3 | Sense: GUACGGUGUCUAGAAUUAAdTdT |
| Antisense: UUAAUUCUAGACACCGUACdTdT | |

**Table S2**. The gRNA sequences for BACH1 knock out

| **Targets** | **gRNA sequence (5′→ 3′)** |
| --- | --- |
| BACH1-1 | FP: AAACACCGCCTGGCCTACGATTCTTGAG |
|  | RP: CTCTAAAACCTCAAGAATCGTAGGCCAGG |
| BACH1-2 | FP: AAACACCGTTACCTTTGAAATCCGACTT |
|  | RP: CTCTAAAACAAGTCGGATTTCAAAGGTAA |

**Table S3**. The primer pairs used for quantitative RT-PCR

| **Genes** | **Nucleotide sequences (5′→ 3′)** |
| --- | --- |
| ACTB | FP: CCAACCGCGAGAAGATGACC |
| BACH1 | RP: GAGTCCATCACGATGCCAGT  FP: GTGGAGCGAGAAGTGGCAGAAC  RP: TACCTAACCACGGACACTCAGACC |
| Caspase3 | FP: ACTGGAATGACATCTCGGTCT |
| BCL2 | RP: ACATCACGCATCAATTCCACA  FP: CTGCACCTGACGCCCTTC  RP: ACACATGACCCCACCGAAC |
| BAX | FP: GACATGTTTTCTGACGGCAAC |
|  | RP: TGTCCAGCCCATGATGGTTC |
| TGFβ1 | FP: GCAACAATTCCTGGCGATACCTC |
|  | RP: AATTTCCCCTCCACGGCTCA |
| Smade2 | FP: GTATTAGTGCCCCGACACACC |
|  | RP: ATTACTCTGTGGCTCAATTCCT |
| Smade3 | FP: CCCCGAAAACACTAACTTCCC |
|  | RP: TTTGGAGAACCTGCGTCCAT |
| Smade4 | FP: AAATATTGTCAGTATGCGTTT |
| PDP1 | RP: TACTTGATGGAGCATTACTCT  FP: GTTGGTGGCTACAAGGTGACTCTG  RP: ATGAGATGGGTTGCTGCGTTCTG |

1.	Kim, D., B. Langmead, and S.L. Salzberg, *HISAT: a fast spliced aligner with low memory requirements.* Nat Methods, 2015. **12**(4): p. 357-60.

2.	Zeiger, W., D. Ito, C. Swetlik, M. Oh-hora, et al., *Stanniocalcin 2 is a negative modulator of store-operated calcium entry.* Mol Cell Biol, 2011. **31**(18): p. 3710-22.

3.	*The Gene Ontology Resource: 20 years and still GOing strong.* Nucleic Acids Res, 2019. **47**(D1): p. D330-d338.

4.	Kanehisa, M. and S. Goto, *KEGG: kyoto encyclopedia of genes and genomes.* Nucleic Acids Res, 2000. **28**(1): p. 27-30.
